# Implementation of Ribo-BiFC method to plant systems using a split mVenus approach

**DOI:** 10.1101/2024.09.12.612679

**Authors:** Karel Raabe, Alena Náprstková, Janto Pieters, Elnura Torutaeva, Veronika Jirásková, Zahra Kahrizi, Christos Michailidis, David Honys

**Author notes:** corresponding authors: KR, DH.

## Abstract

Translation is a fundamental process for every living organism. In plants, the rate of translation is tightly modulated during development and in response to environmental cues. However, it is difficult to measure the actual translation state of the tissues *in vivo*. Here, we report the implementation of an *in vivo* translation marker based on bimolecular fluorescence complementation, the Ribo-BiFC. We combined method originally developed for fruit-fly with an improved low background split-mVenus BiFC system previously described in plants. We labelled *Arabidopsis thaliana* small subunit ribosomal protein (RPS) and large subunit ribosomal protein (RPL) with fragments of the mVenus fluorescent protein. Upon the assembly of the 80S ribosome, the mVenus fragments complemented and were detected by fluorescent microscopy. We show that these recombinant proteins are in close proximity in the tobacco epidermal cells, although the signal is reduced when compared to BiFC signal from known interactors. This Ribo-BiFC method system can be used in stable transgenic lines to enable visualisation of translational rate in plant tissues and could be used to study translation dynamics and its changes during plant development, under abiotic stress or in different genetic backgrounds.

## Background

Translation is one of the fundamental cellular processes, during which proteins are synthesised according to the coding sequence of messenger RNA (mRNA) molecules. In plants, the mechanism of protein synthesis is highly conserved and similar to other eukaryotes (Browning and Bailey-Serres, 2015). Nevertheless, specific ways of regulation emerged in plants, particularly during plant development, in response to abiotic or biotic stresses and response to other environmental stimuli such as light or presence/absence of nutrients (Browning and Bailey-Serres, 2015; Merchante et al., 2017; Urquidi Camacho et al., 2020). The most recognizable component of the translation machinery is the two-subunit ribosome. Ribosomes are large ribonucleoprotein complexes composed of small ribosomal subunit (40S) and large ribosomal subunit (60S). When compared to other eukaryotes, plant ribosomes show some specific differences. For example, the 60S subunit is about 20% smaller than that of mammals (Verschoor and Frank, 1990) and comprises 5S, 5.8S, and 25-26S ribosomal RNAs (rRNA) and approximately 48 ribosomal proteins (RPs) (Barakat et al., 2001). Additionally, plants possess a specific P-protein named P3 (Szick et al., 1998).

Due to translation regulation triggered by quick changes in the environment or connected to developmental progress, the rate of translation changes dynamically in the plant lifespan. Monitoring translational rate has been historically assessed using polysome profiling method, where ribosome subunits, monosomes and polysomes in tissue extract are separated by molecular weight in a sucrose gradient and detected by absorbance profile of the gradient (Mustroph et al., 2009; Mazzoni-Putman and Stepanova, 2018). The polysome/monosome ratio (PM ratio) is generally used to calculate the translation rate and its changes. Although this method is used to analyse the *in vivo* translational state of the tissue, it requires tissue homogenization and extraction. Additionally, several methods based on the use of fluorescently tagged components of the translational machinery exist as well (Mazzoni-Putman and Stepanova, 2018). However, there is no *in vivo* method established in plants that would enable to assess the translation rate of the cell/tissue/organ and its changes using fluorescence microscopy (Mazzoni-Putman and Stepanova, 2018).

A method for *in vivo* fluorescent visualisation of translating ribosomes was described in the *Drosophila melanogaster* system and was used to track assembled, potentially translating ribosomes in Drosophila neurons (Al-Jubran et al., 2013; Singh et al., 2020). This approach is based on ribosomal proteins labelled with a split fluorescent protein, YFP or mVenus, used for Bimolecular fluorescence complementation (BiFC). BiFC is a method canonically used for the verification of protein-protein interactions. Here, however, it is used to label the interaction of 40S and 60S subunits in a way that only assembled 80S ribosomes get the halves of the fluorescent proteins in proximity to the FP complementation and fluorescence signal upon excitation (Kerppola, 2008). Due to the overall conservation of translational machinery in eukaryotes, this method could be possibly implemented in other organisms, including plants. Here we present the implementation of the Ribo-BiFC in plant systems.

We combined the Ribo-BiFC method described in the *Drosophila melanogaster* system (Singh et al., 2020) with the improved low-background split mVenus BiFC system in plants (Gookin and Assmann, 2014). We fused the N-terminal and C-terminal fragments of mVenus to the C-termini of solvent-oriented large and small ribosomal subunit proteins that are near each other in the assembled 80S ribosome. We chose candidates of RPS and RPL proteins based on their position according to the Drosophila Ribo-BiFC and the structure of translating 80S ribosome from *Triticum aestivum* (PDB code: 4V7E) (Armache et al., 2010). Further, as our goal would be implementation to the commonly used model plant, we searched for *Arabidopsis thaliana* orthologues and chose one Arabidopsis paralogue based on conservation and overall expression pattern. We cloned the CDSs of these genes in mVenus-BiFC expression cassettes and screened for mVenus signal in *Nicotiana benthamiana* transient expression system, together with suitable positive and negative controls. We detected fluorescence complementation of tested Ribo-BiFC constructs in the transient expression system, while the negative controls emitted no specific signal. We established that Ribo-BiFC works as a fluorescent translational marker with the potential to visualize translational rate in stable transgenic material *in vivo*, which can be applied for studies of translation dynamics during plant development or translational response to stress conditions.

## Methods

### Plant material and cultivation

*Arabidopsis thaliana* ecotype Columbia-0 inflorescences were used for RNA extraction in order to amplify the CDS sequences of the ribosomal proteins. Seeds of *Arabidopsis thaliana* were surface sterilised (70% ethanol for 1 min, 20% bleach for 10 min, 5x sterile water wash) and germinated on ½ Murashige Skoog media (2.2 g/L MS basal salts; 100 mg/L myo-inositol; 500 mg/L MES; 0.5 mg/L Nicotinic Acid; 0.5 mg/L Pyridoxine·HCl, 1.0 mg/L Thiamine·HCl) pH 5.7 (adjusted with 0.1M KOH solution) to with 0.8% Agar (Sigma). The seeds were stratified below 8°C for 48 hours and germinated at 21°C on long day (16 hr light/8 hr of night) *in vitro*. Seedlings were then transferred to soil (Jiffy tablet) and placed in a growth room (22°C, long day) until flowering. *Nicotiana benthamiana* plants were used for the transient expression experiments. Seeds were germinated in soil for 10 days and transferred to pots, where they were grown at 22°C under white light in a greenhouse. Leaves of juvenile 4–5-week-old plants were used for the leaf infiltration transformation.

### Ribo-BiFC design and cloning

Selected CDS obtained from TAIR (**Supplementary Table 1**) were processed by the GoldenBraid 3.0 domesticator software (https://gbcloning.upv.es/do/domestication/) (Sarrion-Perdigones et al., 2011). The generated oligonucleotides for domestication of the selected CDS were used to domesticate the sequences for the use in the GoldenBraid system. Oligonucleotides for the Ribo-BiFC tags domestication were designed manually to split YFP (NY and CY) and mVenus (NmV and CmV) sequences (**Supplementary Table 2**). YFP was split at amino acid 155, with HA tag sequence fused to the 5’ end of the NY and MYC tag sequence fused to the 5’ end of the CY (**Figure 1B**). mVenus sequence was split at amino acid 210 according to the original publication in plants (Gookin and Assmann, 2014). Additionally, we placed sequence encoding 3xFLAG tag fused to the 5’ end of the small CmV fragment (**Figure 1B**). The NmV has no tag, since it should be big enough protein to be detectable by polyclonal anti-GFP antibody. All oligonucleotides used for domestication are listed in **Supplementary Table 3**. All fragments designed *in silico* were amplified by PCR with Phusion polymerase (Life Technologies) with proofreading activity which was used for fragments amplification. Template for ribosome proteins CDS was the cDNA of *Arabidopsis thaliana* inflorescences obtained using ImProm-IITM Reverse Transcription System (Promega) from RNA isolated from plant tissue using RNeasy plant Mini Kit (Qiagen). The split YFP BiFC fragments were domesticated from the pBiFCt-2in1-CC initial BiFC vector (Grefen and Blatt, 2012). The split mVenus BiFC fragments were domesticated from pUPD2 plasmid containing the mVenus sequence used in (Kubalová et al., 2024) and 3xFLAG from the pICSL50007 plasmid from the Golden Gate plant kit (Engler et al., 2014). PCR-obtained amplicons of the RPL/RPS fragments were cloned into full CDS sequences in pUPD2 vector backbones according to the GoldenBraid restriction-ligation protocol (**Supplemetary Figure 1**) (Sarrion-Perdigones et al., 2011). All cassettes were assembled into full transcription units controlled by a strong viral sporophytic promoter, Cassava vein mosaic virus promoter (pCsvmv) which is comparable to p35S (Verdaguer et al., 1996) and NOSt terminator (**Supplemetary Figure 1A, 1B**). All vectors were cloned in the GoldenBraid cloning system with extended set of assembly vectors (Dusek et al., 2020) (**Supplemetary Figure 1A, 1B**). Chemically competent *E. coli* TOP10a cells were transformed with the ligation reaction by heat shock, plasmid-containing colonies were selected by appropriate antibiotics and blue/white selection. Plasmids were isolated using GeneJET Plasmid Miniprep Kit (Thermo Scientific) and verified by restriction enzyme digest reaction and Sanger sequencing (LightRun - Eurofins).

**Figure 1:**
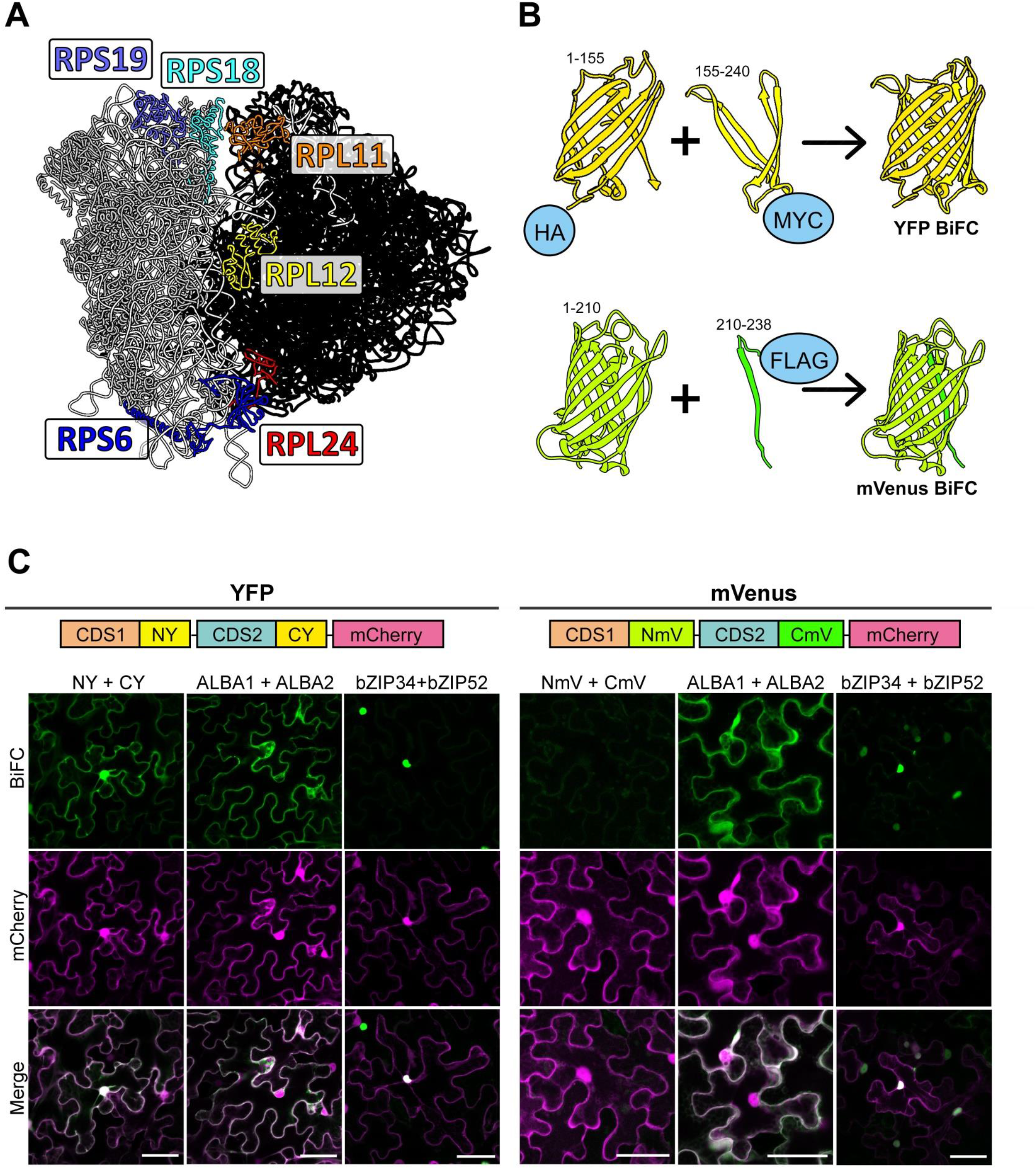
Design of the Ribo-BiFC system and comparison of the split YFP and the split mVenus. **(A**) The structure of the 80S translating ribosome from *Triticum aestivum* (PDB code: 4V7E) visualized in Chimera software. Selected ribosomal proteins are highlighted in various colours. Both rRNA and the rest of RPs are coloured in black or grey for 60S and 40S, respectively. **(B)** Visualization of the BiFC design, depicting the split YFP molecule (upper part) and mVenus (lower part). Numbers indicate amino-acid range of each part of the YFP (NY 1-155; CY 155-240) or mVenus (NmV 1-210; CmV 210-238). Histochemical tags are displayed fused to N-terminal ends of the respective BiFC fragments (HA-YN+MYC-YC and NmV+Flag-CmV). **(C)** Transient coexpression of schematically shown insertion cassettes YFP (left section) or mVenus (right section) BiFC controls with P19 enhancer suppressor are displayed. Detected signal of free FP BiFC fragments is shown in the first columns of each section (NY+CY and NmV+CmV) in transiently transformed tobacco pavement cells. Signal of known interacting partners ALBA1-ALBA2 and bZIP34-bZIP52 was detected and served for the YFP and mVenus BiFC comparison. Data for each sample were obtained by detection of YFP/mVenus signals in green (top row) and a free mCherry signal (middle row) in magenta. Merged channel shows overlays of the BiFC samples with mCherry signal. Scale bar equals to 50 μm.

### *Nicotiana benthamiana* transient assays

*Agrobacterium tumefaciens* competent cells (strain GV3101) were transformed by verified plasmids and plasmid-containing colonies were selected on YEB medium supplemented with gentamicin (50 µg/mL), rifampicin (50 µg/mL), and a vector specific selection agent at 28°C for 48 h. Colonies were inoculated in liquid media and grown overnight at 28°C. Overnight cultures were pelleted by centrifugation (5 min at 1620 g), washed twice, re-suspended, and diluted to an OD^600^ of 0.2 with infiltration medium (10 mM MES pH 5.6, 10 mM MgCl2 and 200 µM acetosyringone). A suspension of *Agrobacterium tumefaciens* cells carrying the plasmid with expression cassette of p19 suppressor of silencing was added in an 1:1 ratio of OD^600^ (Gehl et al., 2009). Mixed suspensions were incubated with moderate shaking for 3 h at room temperature and subsequently injected into the abaxial side of 4-week-old *N. benthamiana* leaves using 1 mL or 2 mL syringe. Two to three days after infiltration, tobacco epidermal cells were analysed microscopically.

### Microscopy and image analysis

Ribo-BiFC of YFP and mVenus signal was localized at subcellular level in pavement cells (*Nicotiana benthamiana*). All microscopic data were obtained by the inverted confocal laser scanning microscope (Axio Observer Z1) Zeiss LSM880 (Carl Zeiss, Jena, Germany) with Plan-Apochromat 20x/0.8 DIC M27 or alpha Plan-Apochromat 100x/1.46 Oil DIC M27, respectively. The fluorophores were visualized by the Argon ion laser 488nm and FS38/GFP BP filter cube (Ex 470/40 Em BP 525/50) (YFP and mVenus) and DPSS laser 561 nm and FS63 HE/mRFP filter cube (Ex BP 572/25 Em BP 629/62) (mCherry) and detected by PMT detector. All acquired images were processed in ZEN blue software (Carl Zeiss, Jena, Germany).

### Structure and gene expression visualization

Structural visualization was performed with Chimera 1.15 software (Pettersen et al., 2004). For ribosomal proteins visualization and positioning within the 80S structure, we used structure of plant 80S translating ribosome from *Triticum aestivum* (PDB code: 4V7E) (Armache et al., 2010). The YFP structure was visualized from template of the crystal structure of eYFP (PDB code: 6VIO) and visualization of the mVenus FP was created by the template structure of mVenus (PDB code: 6SM0). The expression analysis was done with Genevestigator® (https://genevestigator.com/) which uses microarray expression data and showed expression levels for each accession according to the plant anatomy or development. We based our analysis on the Affymetrix GeneChip data visualization using the Development functions. For comparison of Arabidopsis RPs, the protein sequences were downloaded from the TAIR database and aligned with the MUSCLE algorithm.

## Results and Discussion

### Selection of ribosomal proteins for the Ribo-BiFC

This plant-based Ribo-BiFC method for the *in-vivo* visualisation of assembled 80S ribosomes is based on the *Drosophila* Ribo-BiFC approach (Al-Jubran et al., 2013; Singh et al., 2020). There, the tagging of multiple small and large ribosomal protein pairs (combinations of two *Drosophila* ribosomal proteins; RPS18, RPS13, RPL5, RPL11, RPS6 and RPL24) was used. To identify the *Arabidopsis thaliana* orthologues of the RPs in *Drosophila*, we used the assembled 80S ribosome structure from *Triticum aestivum* (Armache et al., 2010) to identify RPs that have similar positions within the ribosome. We then used the *Triticum aestivum* RP amino acid sequences of the proteins for HMMR homology search in *Arabidopsis thaliana* protein database with standard setting (https://www.ebi.ac.uk/Tools/hmmer/). After identifying each corresponding RP in *Arabidopsis*, we looked for its paralogues accession according to the ribosomal protein gene list (Browning and Bailey-Serres, 2015). All the selected RPs had multiple paralogous genes in Arabidopsis (**Supplementary Table 4**). We evaluated the amino acid sequences of the paralogous genes (**Supplementary Table 4**) as well as their expression profiles (**Supplementary Figure 2**). We observed high levels of homology between the paralogs, reaching between 90 % to 100 % sequence identity. The only exception was the RPL24 gene AT2G44860 which shared only around 33 % sequence identity with the other two RPL24 genes. Therefore, the gene was excluded from the final RP selection. Protein sequence comparison gave no indication on one preferable paralogue within the gene family. Therefore, we based our choice on gene expression (**Supplementary Figure 2**). Ribosomal protein genes with overall high expression, and co-expression with other RPs in the whole plant for tagging - RPS6A (At4g31700); RPS18C (At4g09800); RPS19A (At3g02080); RPL11C (At4g18730); RPL12A (At2g37190); RPL24A (At2g36620). All selected RPs were visualized on the plant assembled 80S ribosome structure (**Figure 1A**). According to their position, we identified optimal pairs for Ribo-BiFC as either RPS19 or RPS18 paired with RPL11 and RPS6 paired with RPL24. From this perspective, we hypothesized that RPL12 could also form interaction with RPS18/RPS19 but with lower efficiency than the optimal pairs. This design also allowed tagging RPs that are on opposite sides within the 80S ribosome, potentially representing a non-optimal combination that should possess reduced or entirely lacking the detection of the BiFC signal in the experiments (eg. RPL11 with RPS6). According to this perspective, we hypothesized that when 80S ribosomes are assembled during translation, it would ensure that the selected RPs are brought within close proximity of each other. These RPs pairs tagged with the split YFP/mVenus would serve as a nucleation core for the necessary stabilisation and orientation of the FP parts into the correct conformation and light emission. Confocal microscopy of selected plant tissues could be then performed to visualise the fluorescence intensity that reflects the number of assembled ribosomes.

### Design of the split YFP and mVenus approach for the Ribo-BiFC

Bimolecular fluorescence complementation (BiFC) is a method that was developed to visualise interaction between two proteins when they are in near proximity. A fluorescent protein (FP) is divided into two parts, with each part attached to a different target protein. When the proteins end up in close proximity of around 7 nm (Fan et al., 2008), it results in the FP parts assembly, structure restoration and fluorescence emission upon excitation (Kodama and Hu, 2012; Horstman et al., 2014). The Ribo-BiFC methods developed for *Drosophila melanogaster* showed that neared proteins in the assembled ribosome can also be visualised using the BiFC system. Two fluorescent proteins, YFP and mVenus, were used for our plant adapted method in order to balance the strength of split fluorescent parts interaction and reduction of background noise. The split YFP was divided into NY (amino acid 1-155) and CY (amino acid 155-240), splitting the YFP β-barrel between 7^th^ and 8^th^ β-strand (**Figure 1B**), which is a classical way to split the FP in BiFC methods (Kodama and Hu, 2012). The mVenus sequence was divided into a large NmV (amino acid 1-210) and a small CmV (amino acid 210-238) parts, dividing the β-barrel between the 10^th^ and 11^th^ β-strand. This advanced division has been reported to have high specificity and lower background (Gookin and Assmann, 2014; Ohashi and Mizuno, 2014). We cloned the FP parts as C-terminal fusions to the CDS sequences. Additionally, we enriched the parts by histochemical tags between CDS and FPs that enables histochemical protein detection easier and serves also as a linker sequences (ct-HA-NY, ct-MYC-CY, ct-Flag-CmV).

### Assembly of the Ribo-BiFC expression vector

The CDS sequences of the selected genes encoding ribosomal proteins were isolated and domesticated to the GoldenBraid cloning system with sequence tags suitable for C-terminal fusions. We chose strong viral promoter of Cassava vein mosaic virus (pCsVMV) to drive the expression of the Ribo-BiFC, BiFC and control constructs, since it is a promoter active in both *N. benthamiana* transient expression system, as well as in *Arabidopsis thaliana* stable lines (for possible use of the same expression cassettes in stable lines). We prepared single full transcription units (TUs) in expression clones and continued to the destination vector by their combination according to the scheme (**Supplementary Figure 1**). The destination backbone is the GoldenBraid 3.0 binary vector pDGB3 Ω2 and the inserts were assembled in one restriction/ligation reaction, using the α11 to α14 plus α2 system (Dusek et al., 2020). The α11 position contained the BiFC expression cassette with FP-Nt fragment (pCsVMV::CDS1-NY/NmV::NOSt), while the α12 position contained the BiFC expression cassette with FP-Ct fragment (pCsVMV::CDS2-CY/CmV::NOSt). Then, α13 and α14 positions were filled with 35bp stuffer (sf) sequence and α2 position contained the cassette expressing free mCherry (pCsVMV::mCherry::NOSt) that served for successfully transformed *N. benthamiana* cells identification. We also cloned the NY, CY, NmV and CmV parts as CDS sequences to express them freely in the *Nicotiana benthamiana* cells as one set of negative controls. In summary, the whole Ribo-BiFC is transferred to plants using one vector.

### Split mVenus BiFC shows lower background noise than split YFP

The transient expression of the BiFC controls in *N. benthamiana* showed clear difference in the split YFP and mVenus molecular tags (**Figure 1C**). While the expression of the commonly used free YFP parts (NY+CY) showed non-specific complementation and strong signal emission in the cytoplasm and nucleus, the free parts of mVenus210 (NmV+CmV) revealed lower background signal, almost comparable with the set of negative controls of P19-transformed, mock-transformed (infiltration media only) and non-transformed cells (**Supplementary Figure 3**). We also tested known interacting partners, ALBA1+ALBA2 (Yuan et al., 2019), as positive controls with strong signal in cytoplasm and two dimerizing members of bZIP family transcription factors known to form dimers in nucleus, bZIP34+bZIP52 (Gibalová et al., 2017) with the same result in both BiFC approaches (**Figure 1C**). Moreover, the identical complemented signal pattern corresponds with our previous localization of ALBA family proteins in cytoplasm (Náprstková et al., 2021; Tong et al., 2022) and bZIP proteins in nucleus (Gibalová et al., 2017). Altogether, we concluded that the mVenus BiFC is superior to the canonical YFP tag. Although we tested most of the Ribo-BiFC combinations with both YFP and mVenus system with similar results, we only show here the novel mVenus Ribo-BiFC.

### Ribo-BiFC in Nicotiana benthamiana pavement cells

The selected Ribo-BiFC pairs signal strength corresponded to the position of the RP pair on the ribosome (**Figure 2**). We detected prominent cytoplasmic mVenus signal in promising Ribo-BiFC pairs RPS18+RPL11, RPS19+RPL11 and moderate signal emission of RPS6+RPL24. In comparison, we observed weaker or no signal in the sub-optimal Ribo-BiFC pairs, RPS18+RPL12 and RPS19+RPL12. Interestingly, RPLx-NmV+RPSx-CmV pairs revealed decreased signal intensity emission in the cytoplasm; the most differences were discovered in between RPS18+RPL11 and RPL11+RPS18 reciprocal pairs as well as in RPS6+RPL24 and RPL24+RPS6. The mVenus signal was detected in nucleoli, but also in nuclei and cytoplasm. The strong signal in nucleoli might be due to RPs accumulation in this compartment where ribosomes are assembled, as the high number of ribosomal proteins fusions produced under the viral promoter is excessive to the number of ribosomes being assembled in the nucleolus. We tested the localization of selected RPs by expression cassettes, where the RPs are fused to full mVenus on their C-terminal (**Supplementary Figure 4**). These constructs showed a similar pattern of localization. The mVenus signal was detected in the cytoplasm, nucleus and strongly in the nucleolus. Additionally, similar localization was reported in other ribosomal protein studies in Arabidopsis (Yao et al., 2008) or rice (Zheng et al., 2016). To conclude, the revealed Ribo-BiFC signal pattern matches the localization of the RPs. Our hypothesis is that the signal presence in nucleolus is an artefact due to the overexpression rather than a visualization of assembled ribosomes.

**Figure 2:**
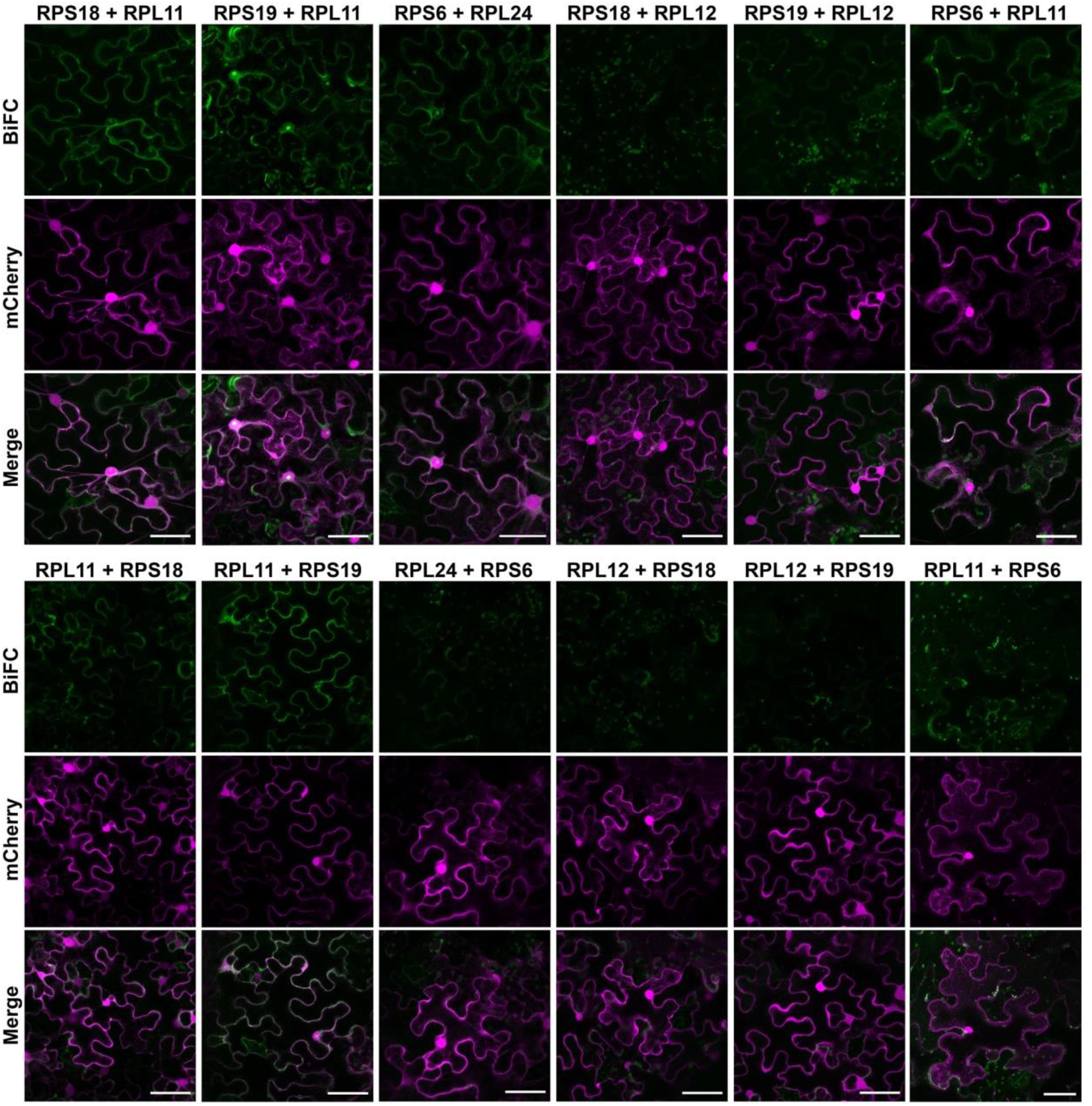
Ribo-BiFC combinations in *Nicotiana benthamiana*. Transient tobacco assay of selected Ribo-BiFC combinations revealed variability in detected signal intensity. Each combination lists two RP fusions, where the first RP is fused to NmV and the second RP is fused to CmV. The upper section shows combinations RPS-NmV + RPL-CmV. The lower section shows combinations RPL-NmV + RPS-CmV. Results of the signal detection are shown in the green channel for mVenus BiFC. Free mCherry control of the pavement cells transient transformation is in the middle (shown in magenta) and the overlay of the channels is displayed in the third row. All transcription units are driven by pCsVMV promoter and acquired data were processed the same way. Scale bars equal to 50 μm.

In comparison to the usual signal strength of standard BiFC, for example ALBA1-ALBA2, the Ribo-BiFC mVenus signal was decreased. Given the fact the only assembled ribosomes with both protein fusions integrated can complement the FP emission, the reduced signal strength suggests tracking dynamic structures, the assembled ribosomes.

To exclude the possibility of non-specific complementation, we chose to use the RPL11 and RPS19 proteins to be tested in a set of BiFC controls (**Supplementary Figure 3**). We tested combinations of the free NmV and CmV in a pair with either RPL11 or RPS19 tagged with the complementary mVenus fragment. We detected no mVenus signal in these control BiFC combinations, with exception of a very low signal in the nucleolus in some transformed cells. We additionally tested bZIP52 with RPL11 or RPS19 in similar way, with weak background signal in nucleus, in case of the combination bZIP52 + RPS19. Although our results indicate non-specific signal also in the mVenus BiFC, the detected background signal is far below YFP BiFC background intensity. Moreover, the signal in the optimal Ribo-BiFC combinations was stronger than in negative controls and the suboptimal Ribo-BiFC pairs.

## Conclusions

Here, we describe the implementation of a method into plant systems, the Ribo-BiFC. This approach is based on the previously reported *Drosophila melanogaster* Ribo-BiFC method, which involves tagging small and large ribosomal proteins with BiFC fragments to visualise their proximity when ribosomes assemble during translation. To improve the signal specificity, we have designed a novel BiFC using mVenus FP split in a way that reduces the non-specific BiFC signal. We obtained the split mVenus fragments in the GoldenBraid system that enables assembly of all Ribo-BiFC parts into one vector. Further, we tested several candidate pairs of Arabidopsis thaliana ribosomal proteins in Tobacco transient assays. We were able to detect stronger mVenus signal in Ribo-BiFC pavement cells than the background of negative controls despite of its reduction compared to the known interactors. The Ribo-BiFC revealed a signal localized in the nucleolus, which is most probably an overexpression artefact. Furthermore, the signal pattern did not differ from single RP overexpression in the same transient system.

Finally, this study introduces the Ribo-BiFC implementation in plants. We aim to produce on stable Ribo-BiFC lines of *Arabidopsis thaliana*. This establishment is crucial for advanced experiments and treatments that would show Ribo-BiFC fluorescence changes together with changes in translation rate measured by polysome/monosome ratio calculation. Such treatments could include environmental stresses and translation-drug treatments. Since a search for most stress-free conditions does not correspond with the strong viral promoter that causes free RPs accumulation, we suggest achieving native expression of the RPs by either UBQ10, native promoter of the used RPs, or their combination that could even reduce the rate of the possible T-DNA silencing.

Consequently, Ribo-BiFC stable line would permit non-invasive assessment of the translational rate of any given tissue or organ in different genetic backgrounds or under various environmental or biological conditions. Considering the conserved structure of the translational machinery, this approach could be introduced in other plant species as well and bring new insights into one of the most dynamic and essential molecular processes.

## Acknowledgement

We acknowledge the Imaging Facility of the Institute of Experimental Botany AS CR supported by the MEYS CR (LM2023050 Czech-BioImaging), the Czech Academy of Sciences and IEB AS CR.

