## Supplementary_material for "Implementation of Ribo-BiFC method to plant systems using a split mVenus approach"

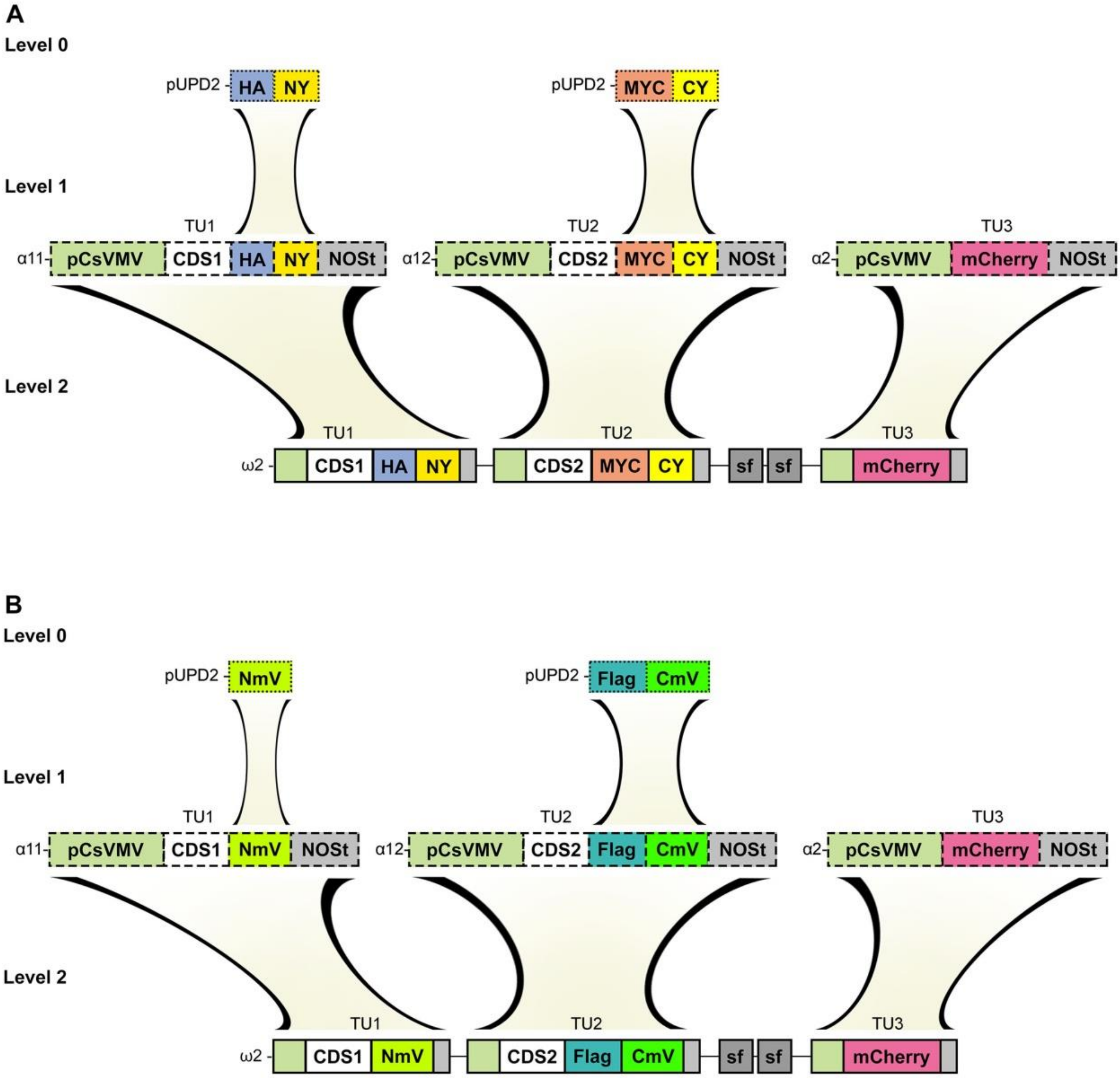
A
Level 0
pUPD2
HA
NY
pUPD2
MYC
CY
Level 1
TU1
TU2
TU3
α11
pCsVMV
CDS1
HA
NY
NOSt
α12
pCsVMV
CDS2
MYC
CY
NOSt
α2
pCsVMV
mCherry
NOSt
Level 2
TU1
TU2
TU3
ω2
CDS1
HA
NY
CDS2
MYC
CY
sf
sf
mCherry
B
Level 0
pUPD2
NmV
pUPD2
Flag
CmV
Level 1
TU1
TU2
TU3
α11
pCsVMV
CDS1
NmV
NOSt
α12
pCsVMV
CDS2
Flag
CmV
NOSt
α2
pCsVMV
mCherry
NOSt
Level 2
TU1
TU2
TU3
ω2
CDS1
NmV
CDS2
Flag
CmV
sf
sf
mCherry

**Supplementary Figure 1: Schematic overview of the destination vector assembly**

The individual steps were based on the GoldenBraid 3.0 cloning. (A) At the Level 0, fragments of the NY and CY were domesticated from the pBiFCt-2in1-CC vector and N-terminally fused with HA or MYC tags domesticated from MoClo. The selected RPs coding sequences were domesticated according to Sarrion-Perdigones et al., 2011. Next, the transcription units were assembled at Level 1 and combined to the destination vectors at Level 2. (B) Fragments of NmV and CmV were domesticated from full mVenus sequence obtained from Kubalová et al., 2024. While NmV size is sufficient for the GFP antibody recognition, CmV was N-terminally fused with Flag tag. Both domesticated split mVenus parts were canonically implemented into Level 1 and Level 2 GoldenBraid cloning.

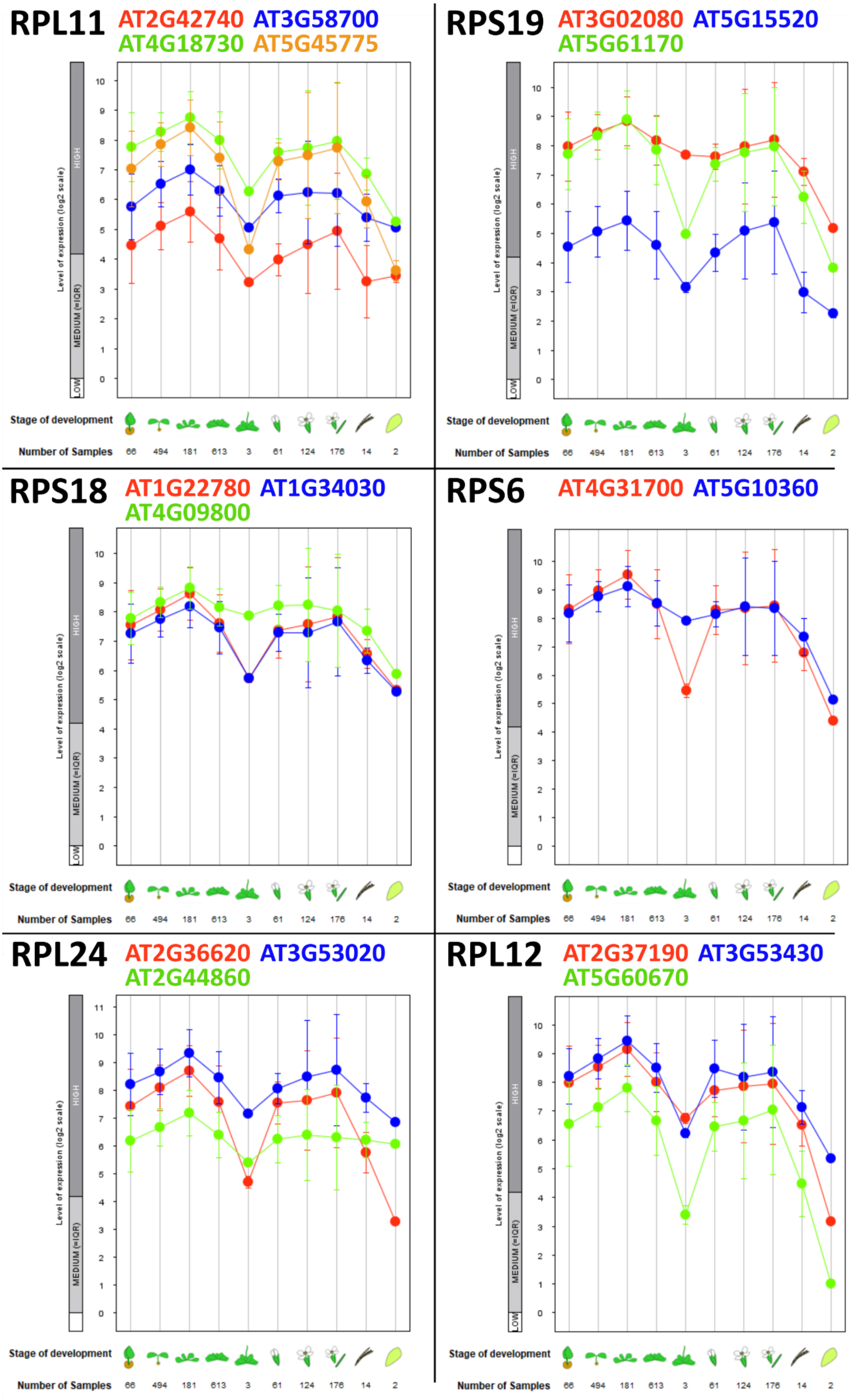

RPL11 AT2G42740 AT3G58700 AT4G18730 AT5G45775
RPS19 AT3G02080 AT5G15520 AT5G61170
RPS18 AT1G22780 AT1G34030 AT4G09800
RPS6 AT4G31700 AT5G10360
RPL24 AT2G36620 AT3G53020 AT2G44860
RPL12 AT2G37190 AT3G53430 AT5G60670
Level of expression (log2 scale)
HIGH
MEDIUM (=IQR)
LOW
Stage of development
Number of Samples
66 494 181 613 3 61 124 176 14 2

**Supplementary Figure 2: RP paralogues expression profiles**

The expression profiles for the Arabidopsis thaliana RPs paralogue genes are displayed according to Affymetrix GeneChip data and were generated using the Development functions of Genevestigator. The expression profiles are shown for RPL11 (upper left), RPS18 (middle left), RPL24 (lower left), RPS19 (upper right), RPS6 (middle right), and RPL12 (lower right). Data from ATH arrays are shown in scatter-plot diagrams. The x-axis represents the following developmental stages, from left to right: germinating seed, seedling, young rosette, developed rosette, bolting, young flower, developed flower, flowers and siliques, mature siliques, and senescent leaves. For each data point, the number of samples is indicated. The values in the plots are the mean values, and the error bars show standard errors.

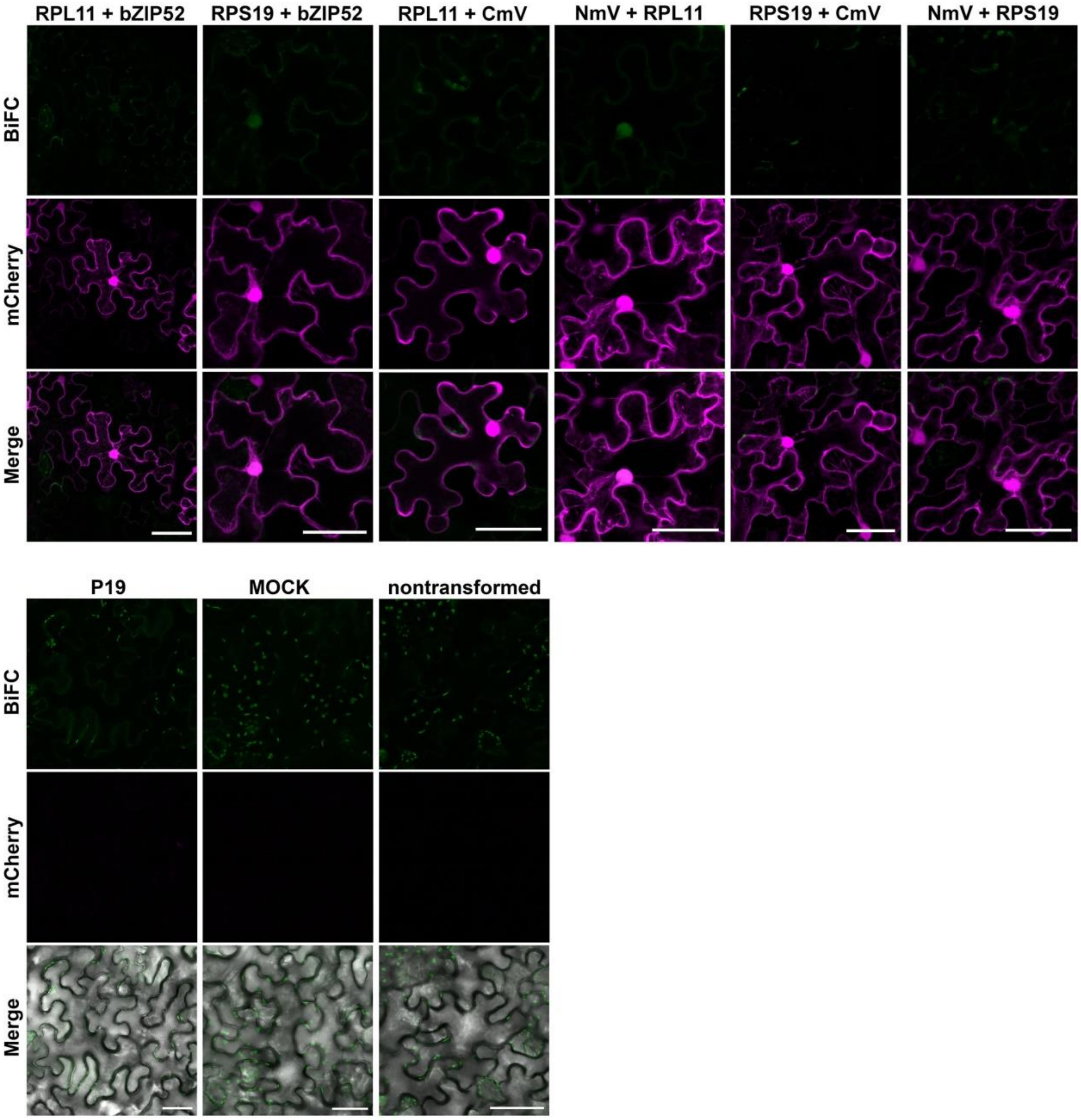
RPL11 + bZIP52
RPS19 + bZIP52
RPL11 + CmV
NmV + RPL11
RPS19 + CmV
NmV + RPS19
BiFC
mCherry
Merge
P19
MOCK
nontransformed
BiFC
mCherry
Merge

**Supplementary Figure 3: mVenus Ribo-BiFC tested negative controls in pavement cells of *N. benthamiana* transiently transformed leaves**

For transient tobacco assay non-interacting partners were selected (upper row). The samples of negative controls include protein-protein combinations, RPs and bZIP52 fused to NmV or CmV, followed by protein-split mVenus pairs, RPs with free BiFC complementing fragments (free NmV or free CmV). Constructs are driven by pCsvmv promoter, coexpressed with free mCherry and coinfiltrated with P19 supressor. The list of negative controls includes only P19 and infiltration media (MOCK) samples completed by nontransformed pavement cells autofluorescence (bottom row). For each sample, the Ribo-BiFC detection channel is shown in green, transformation control of free mCherry channel is shown in the magenta. The merged image combines both. Scale bars equal to 50 µm.

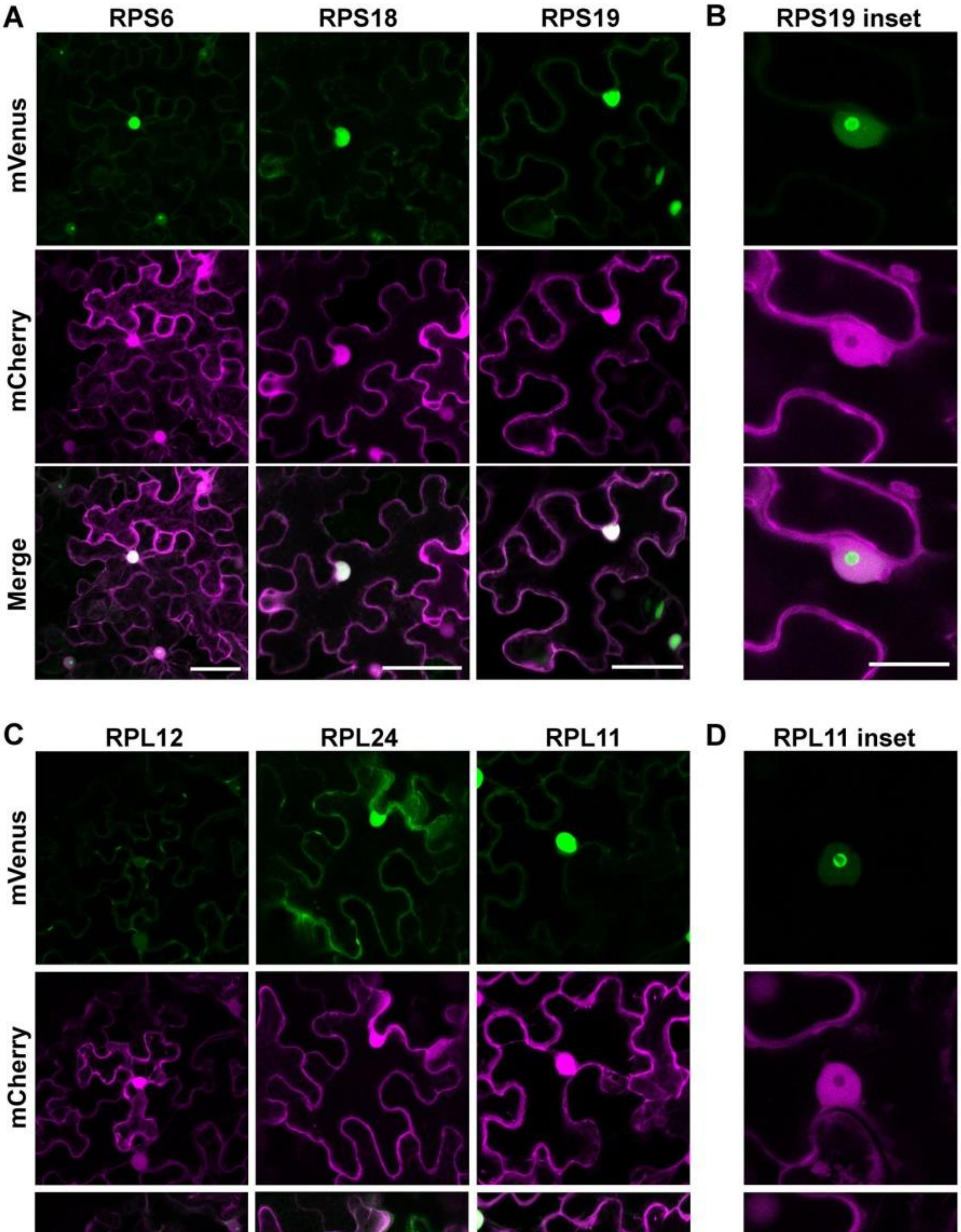
A
RPS6
RPS18
RPS19
B
RPS19 inset
mVenus
mCherry
Merge
C
RPL12
RPL24
RPL11
D
RPL11 inset
mVenus
mCherry
Merge

**Supplementary Figure 4: Subcellular localization of mVenus tagged proteins in *N. benthamiana* transiently transformed pavement cells.**

Selected proteins were fused to full mVenus sequence and served as controls to Ribo-BiFC experiments. (A, C) For each sample, the protein-specific localization is displayed in the green channel (mVenus) and transformation control in magenta (free mCherry). The merge of the channels is shown at the bottom. Scale bar equals to 50 μm. (B, D) Nuclear and possible nucleolar localization of the RPs is represented by RPS19 and RPL11 insets in the green channel. All protein fusions are driven by pCsVMV promoter. Scale bars equal to 20 μm.

### Supplementary Table 1: Sequences of RPs for domestication

| Domesticated part | CDS sequence (Start to stop codon) |
|---|---|
| **RPL11-At4g18730_CDS** | ATGGCATCGGAGAAGAAGCTCTCGAACCCTATGAGGGATATTAAGGTCCAGAAGCTAGTTCTTAACATCTCTGTTGGTGAG<br>AGTGGTGATCGTCTCACTCGTGCCTCCAAGGTGTTGGAACAGCTCAGTGGTCAGACTCCTGTCTTCTCTAAGGCGAGGTAC<br>ACTGTGAGGTCTTTCGGTATCAGGCGTAATGAAAAGATTGCGTGCTATGTCACCGTGAGAGGTGAGAAGGCAATGCAGCT<br>TCTTGAGAGTGGCTTGAAAGTGAAGGAATACGAGCTGTTGAGGAGGAACTTCAGTGACACTGGCTGCTTCGGATTCGGTA<br>TCCAGGAGCACATTGATCTTGGAATCAAGTATGATCCTTCTACCGGTATCTACGGTATGGATTTCTACGTTGTTCTTGAACGT<br>CCAGGATACCGTGTGGCCCGTCGCCGTAGATGCAAGACTCGCGTTGGTATTCAACATAGAGTTACCAAGGATGATGCCATG<br>AAGTGGTTCCAAGTTAAGTATGAAGGAGTTATCCTCAACAAGTCTCAGAACATCACTGGTTGA |
| **RPL12-At2g37190_CDS** | ATGCCGCCAAAGTTGGATCCGAGTCAGATCGTTGACGTGTACGTCCGAGTAACCGGGGGAGAAGTCGGAGCAGCGAGTTC<br>TCTAGCTCCCAAAATCGGTCCGCTAGGTCTCGCACCAAAGAAGATCGGAGAAGATATCGCGAAAGAAACCGCAAAGGAAT<br>GGAAAGGATTACGTGTAACCGTCAAGCTCACAGTGCAGAATCGTCAGGCGAAAGTGACGGTGGTTCCATCAGCAGCAGCA<br>CTCGTCATCAAGGCGTTGAAAGAGCCGGAGAGAGATAGGAAGAAGGTGAAGAACATTAAGCATAACGGGAACATCTCGT<br>TTGATGATGTGACTGAGATTGCTAGGATCATGAGACCTAGATCTATTGCTAAGGAATTGAGTGGAACAGTGAGGGAGATTC<br>TGGGAACTTGTGTGTCTGTTGGTTGTACTGTGGATGGGAAAGATCCTAAAGATATTCAACAGGAGATTCAAGATGGTGAA<br>GTTGAGATTCCTGAGAACTGA |
| **RPL24-At2g36620_CDS** | ATGGTTCTCAAGACTGAGCTTTGCCGATTCAGTGGCCAGAAAATTTACCCTGGTAGAGGGATCAGATTTATCCGATCGGAC<br>TCTCAGGTGTTTTTGTTTCTCAACTCCAAATGTAAGAGGTATTTCCACAACAAGTTGAAGCCATCTAAGCTTTGCTGGACTGC<br>TATGTACCGAAAGCAGCACAAGAAGGACGCAGCACAAGAGGCTGTGAAGAGAAGGAGACGTGCAACTAAGAAGCCTTAC<br>TCAAGGTCGATTGTCGGTGCTACTTTGGAGGTTATTCAGAAGAAGCGAGCAGAGAAGCCTGAAGTTCGTGATGCCGCTAG<br>AGAAGCTGCCCTACGTGAGATCAAGGAGAGAATCAAGAAGACCAAGGACGAGAAGAAGGCAAAGAAGGTCGAGTATGC<br>ATCAAAGCAACAGAAGTCACAAGTGAAGGGAAATATCCCCAAGAGTGCTGCACCCAAGGCTGCTAAGATGGGTGGTGGTG<br>GAGGCAGACGTTGA |
| **RPS6-At4g31700_CDS** | ATGAAGTTCAACGTTGCGAATCCAACTACTGGATGCCAGAAGAAGCTCGAGATCGACGATGACCAGAAACTACGTGCGTTT<br>TACGACAAGAGAATCTCTCAAGAAGTCAGTGGAGATGCTTTGGGCGAGGAGTTCAAAGGATACGTTTTCAAGATCAAGGG<br>TGGTTGCGATAAGCAAGGTTTCCCAATGAAGCAGGGAGTTTTGACTCCAGGCCGTGTTCGCCTTTTGCTTCACCGAGGAAC<br>TCCTTGCTTCAGAGGACATGGAAGGAGAACTGGTGAGAGGAGAAGAAAGTCTGTTCGTGGTTGCATTGTGAGCCCTGATC<br>TCTCTGTTCTGAACCTTGTCATTGTGAAGAAGGGTGAGAACGATCTTCCTGGGCTTACCGATACTGAGAAGCCAAGAATGA<br>GAGGACCAAAGAGAGCCTCCAAGATCCGTAAACTGTTTAACCTCAAGAAGGAAGATGATGTCAGGACCTATGTCAACACTT<br>ACCGCCGCAAGTTCACAAACAAGAAGGGCAAGGAAGTTAGCAAAGCCCCTAAGATCCAGAGGCTTGTGACCCCATTGACT<br>CTTCAGAGGAAGAGAGCTAGAATTGCTGACAAGAAGAAGAAAATTGCTAAGGCTAATTCTGATGCTGCTGATTACCAGAA<br>GCTTCTCGCCTCGAGGTTGAAGGAACAGCGTGACAGGAGGAGTGAGAGTTTGGCAAAGAAGAGGTCGAGACTCTCTTCTG<br>CTGCTGCCAAGCCCTCTGTCACAGCTTAA |
| **RPS18-At4g09800_CDS** | ATGTCTCTGGTTGCAAATGAGGAGTTTCAACACATTCTTCGTGTGTTGAATACTAATGTTGATGGTAAGCAGAAGATTATGT<br>TTGCCCTTACCTCTATCAAAGGTATTGGTAGGCGATTGGCTAACATTGTCTGCAAGAAGGCTGATGTCGACATGAACAAAA<br>GGGCTGGTGAGTTATCTGCTGCTGAGATTGATAACCTCATGACAATCGTTGCAAACCCACGTCAGTTCAAGATCCCAGACTG<br>GTTCTTGAACAGGCAGAAGGATTACAAAGATGGCAAGTATTCTCAAGTTGTCTCCAATGCTCTTGACATGAAGCTGAGAGA<br>TGATCTTGAACGTCTCAAGAAGATCAGAAACCACCGTGGTTTGAGGCATTACTGGGGTCTCCGTGTCAGAGGACAACACAC<br>CAAGACTACTGGTCGCAGAGGAAAGACTGTTGGTGTCTCAAAGAAGCGTTAA |
| **RPS19-At3g02080_CDS** | ATGGCAACTGGTAAAACTGTGAAAGACGTCTCGCCTCATGACTTCGTCAAGGCTTATGCTTCTCATCTCAAGCGATCTGGCA<br>AGATCGAGCTTCCCACATGGACAGACATTGTGAAGACCGGAAAGTTGAAGGAGCTTGCACCATATGATCCTGATTGGTACT<br>ACATCAGAGCTGCATCTATGGCAAGGAAGGTTTACCTGAGGGGAGGACTTGGTGTTGGTGCTTTCCGTAGAATCTATGGTG<br>GAAGCAAGAGGAACGGTAGTCGCCCACCTCACTTTTGCAAAAGCAGTGGTGGTATTGCCCGTCACATCCTCCAACAGCTGG<br>AGACAATGAACATTGTTGAGCTCGACACCAAAGGAGGAAGAAGGATCACTTCCAGTGGCCAAAGGGATTTGGACCAGGTT<br>GCTGGCCGTATTGCAGTTGAACCTTGA |

### Supplementary Table 2: Sequences of BiFC fragments

| Domesticated part | CDS sequence |
|---|---|
| NmV<br>(mVenus-Nt) | GGAGGAGGAAGCAAAGGCGAGGAACTTTTTACCGGGGTTGTTCCCATATTGGTGGAACTAGACGGAGATGTAAATGGCC<br>ACAAGTTCTCAGTTTCCGGTGAGGGTGAGGGTGACGCGACATATGGTAAACTGACCCTGAAATTAATTTGTACCACGGGT<br>AAGTTACCGGTCCCCTGGCCGACTTTGGTCACCACGCTTGGATATGGCGTGCAATGTTTTGCGCGTTATCCAGATCATATG<br>AAGCAACATGATTTCTTTAAGTCTGCAATGCCCGAAGGATATGTGCAGGAGAGAACTATCTTCTTTAAAGACGACGGAAA<br>CTATAAAACAAGGGCCGAGGTTAAGTTCGAAGGAGATACCCTAGTTAACAGAATCGAGTTGAAGGGGATAGATTTTAAG<br>GAGGACGGCAATATCTTAGGTCACAAATTGGAATATAATTACAATAGCCATAACGTATATATCACCGCCGACAAACAAAAA<br>AACGGTATAAAAGCTAATTTCAAAATCAGGCACAACATCGAGGACGGCGGCGTACAGTTGGCTGATCATTATCAACAAAA<br>TACCCCTATAGGTGACGGACCGGTCTTGCTTCCGGACAATCACTATCTCAGTTACCAATCAAAACTCAGCAAAGACTAA |
| CmV<br>(FLAG-mVenus-Ct) | ATGGATTATAAGGACCATGACGGAGACTATAAGGACCATGACCTCGACGCTGCAGCAGCGGATTATAAGGACGATGACG<br>ATAAGCAGCTTCGAGTACCACTCGAGGCCATGCCCAATGAGAAAAGAGATCACATGGTTCTTTTAGAGTTCGTAACCGCG<br>GCTGGCATAACACACGGCATGGACGAGCTATACAAGTAA |
| NY<br>(HA-YFP-Nt) | ATGTATCCTTATGATGTTCCTGATTATGCTACTAGTGGAGGTGGATCTGGAGGTGGAAGTAGAATGGTGAGCAAGGGCGA<br>GGAGCTGTTCACCGGGGTGGTGCCCATCCTGGTCGAGCTGGACGGCGACGTAAACGGCCACAAGTTCAGCGTGTCCGGC<br>GAGGGCGAGGGCGATGCCACCTACGGCAAGCTGACCCTGAAGTTCATCTGCACCACCGGCAAGCTGCCCGTGCCCTGGCC<br>CACCCTCGTGACCACCTTCGGCTACGGCCTGCAGTGCTTCGCCAGGTACCCCGACCACATGAAGCAGCACGACTTCTTCAA<br>GTCCGCCATGCCCGAAGGCTACGTCCAGGAGAGAACCATCTTCTTCAAGGACGACGGCAACTACAAGACCAGGGCCGAG<br>GTGAAGTTCGAGGGCGACACCCTGGTGAACAGAATCGAGCTGAAGGGCATCGACTTCAAGGAGGACGGCAACATCCTGG<br>GGCACAAGCTGGAGTACAACTACAACAGCCACAACGTCTATATCATGGCCTGA |
| CY<br>(MYC-YFP-Ct) | ATGGAACAGAAGCTTATCTCAGAGGAGGACCTGGCCGGCACTAGTGGAGGTGGATCTGGAGGTGGAAGTATGGACAAG<br>CAGAAGAACGGCATCAAGGTGAACTTCAAGATCAGGCACAACATCGAGGACGGCAGCGTGCAGCTCGCCGACCACTACC<br>AGCAGAACACCCCCATCGGCGACGGCCCCGTGCTGCTGCCCGACAACCACTACCTGAGCTACCAGTCCGCCCTGAGCAAA<br>GACCCCAACGAGAAGAGAGATCACATGGTCCTGCTGGAGTTCGTGACCGCCGCCGGGATCACTCTCGGCATGGACGAGC<br>TGTACAAGTAG |

### Supplementary Table 3: Oligonucleotide sequences for domestication of RPs or BiFC

| Oligonucleotide name | Oligonucleotide sequence (5´-> 3´) |
|---|---|
| RPL11_AATG_dom_F1 | GCGCCGTCTCGCTCGAATGGCATCGGAGAAGAAGCT |
| RPL11_dom_R1 | GCGCCGTCTCGTAAGACGATCACCACTCTCAC |
| RPL11_dom_F2 | GCGCCGTCTCGCTTACTCGTGCCTCCAAGGT |
| RPL11_TTCG_dom_R2 | GCGCCGTCTCGCTCACGAACCACCAGTGATGTTCTGAGACTT |
| RPL12_AATG_dom_F1 | GCGCCGTCTCGCTCGAATGCCGCCAAAGTTGGATCC |
| RPL12_dom_R1 | GCGCCGTCTCGCAAGACCTAGCGGACCGATT |
| RPL12_dom_F2 | GCGCCGTCTCGCTTGCACCAAAGAAGATCGG |
| RPL12_dom_R2 | GCGCCGTCTCGGCCTCATGATCCTAGCAATC |
| RPL12_dom_F3 | GCGCCGTCTCGAGGCCTAGATCTATTGCTAAG |
| RPL12_dom_TTCG_R3 | GCGCCGTCTCGCTCACGAACCGTTCTCAGGAATCTCAACTTC |
| RPL24_AATG_dom_F1 | GCGCCGTCTCGCTCGAATGGTTCTCAAGACTGAGCTT |
| RPL24_dom_R1 | GCGCCGTCTCGGCCTCCTTCTCTTCACAGCC |
| RPL24_dom_F2 | GCGCCGTCTCGAGGCGTGCAACTAAGAAGCC |
| RPL24_TTCG_dom_R2 | GCGCCGTCTCGCTCACGAACCACGTCTGCCTCCACCACCAC |
| RPS6_AATG_dom_F1 | GCGCCGTCTCGCTCGAATGAAGTTCAACGTTGCGAATC |
| RPS6_TTCG_dom_R1 | GCGCCGTCTCGCTCACGAACCAGCTGTGACAGAGGGCTTGG |
| RPS18_AATG_dom_F1 | GCGCCGTCTCGCTCGAATGTCTCTGGTTGCAAATGAG |
| RPS18_dom_R1 | GCGCCGTCTCGTAAGACGTTCAAGATCATCTCTC |
| RPS18_dom_F2 | GCGCCGTCTCGCTTAAGAAGATCAGAAACCACCGTGGTTTGAGGCATTACTGGGGTCTTCGTGTCAGA |
| RPS18_TTCG_dom_R2 | GCGCCGTCTCGCTCACGAACCACGCTTCTTTGAGACACCAAC |
| RPS19_AATG_dom_F1 | GCGCCGTCTCGCTCGAATGGCAACTGGTAAAACTGTGAAAGACGTTTCGCCTCAT |
| RPS19_TTCG_dom_R1 | GCGCCGTCTCGCTCACGAACCAGGTTCAACTGCAATACGGC |
| YFP-Nt_fusion_F1 | GCGCCGTCTCGCTCGTTCGATGTATCCTTATGATGTTCCTGAT |
| YFP-Nt_fusion_R1 | GCGCCGTCTCGCTCAAAGCTCAGGCCATGATATAGACGTT |
| YFP-Nt_CDS_F1 | GCGCCGTCTCGCTCGAATGTATCCTTATGATGTTCCTGATTATG |
| YFP-Nt_CDS_R1 | GCGCCGTCTCGCTCAAAGCTCAGGCCATGATATAGACGTT |
| YFP-Ct_fusion_F1 | GCGCCGTCTCGCTCGTTCGATGGAACAGAAGCTTATCTCAG |
| YFP-Ct_fusion_R1 | GCGCCGTCTCGCTCAAAGCCTACTTGTACAGCTCGTCCA |
| YFP-Ct_CDS_F1 | GCGCCGTCTCGCTCGAATGGAACAGAAGCTTATCTCAGAG |
| YFP-Ct_CDS_R1 | GCGCCGTCTCGCTCAAAGCCTACTTGTACAGCTCGTCCA |
| mVenus-Nt_fusion_F1 | GCGCCGTCTCGCTCGTTCGGGAGGAGGAAT |
| mVenus-Nt_fusion_R1 | GCGCCGTCTCGCTCAAAGCTTAGTCTTTGCTGAGTTTTGATTG |
| mVenus-Nt_CDS_F1 | GCGCCGTCTCGCTCGAATGAGCAAAGGCGAGGAA |
| mVenus-Nt_CDS_R1 | GCGCCGTCTCGCTCAAAGCTTAGTCTTTGCTGAGTTTTGATTG |
| mVenus-Ct_fusion_F1 | GCGCCGTCTCGCTCGTTCGGATTATAAGGACCATGACGGA |
| mVenus-Ct_fusion_R1 | GCGCCGTCTCGCGAGTGGTACTCGAAGCTGCTTATCGTC |
| mVenus-Ct_fusion_F2 | GCGCCGTCTCGCTCGAGGCCATGCCCAATGAGAAAAGAGATC |
| mVenus-Ct_fusion_R2 | GCGCCGTCTCGCTCAAAGCTTACTTGTATAGCTCGTC |
| mVenus-Ct_CDS_F1 | GCGCCGTCTCGCTCGAATGGATTATAAGGACCATGACGGA |
| mVenus-Ct_CDS_R1 | GCGCCGTCTCGCGAGTGGTACTCGAAGCTGCTTATCGTC |
| mVenus-Ct_CDS_F2 | GCGCCGTCTCGCTCGAGGCCATGCCCAATGAGAAAAGAGATC |
| mVenus-Ct_CDS_R2 | GCGCCGTCTCGCTCAAAGCTTACTTGTATAGCTCGTC |
| His_fusion_F1 | GCGCCGTCTCGCTCGTTCGGGTTCCGGAAGAGGATCG |
| His_fusion_R1 | GCGCCGTCTCGCTCAAAGCTCACTTGTCATCGTCATCC |

**Supplementary Table 4: RP paralogues sequence identity and paralogue selection**

Tables of sequence identity matrix for the Arabidopsis thaliana RP paralogue genes are shown in upper part. Protein sequences were obtained from the TAIR database at the respective AGI code and aligned with the MUSCLE algorithm. The colour of the box represents the percentage of sequence identity (gradient from low red to high green identity). Selected genes for cloning are highlighted in red. Lower part summarizes the selection of RPs. In the left column, a RP chosen in *Drosophila melanogaster* publication is listed. In the middle column, the RP with a similar position within the 80S *Triticum aestivum* model is listed. The right column lists all paralogue genes in *Arabidopsis*, where the chosen paralogue used in Ribo-BiFC is underlined.

| RPS6 | AT4G31700 | AT5G10360 |
|---|---|---|
| AT4G31700 | 100 | 95.18 |
| AT5G10360 | 95.18 | 100 |

| RPL11 | AT2G42740 | AT3G58700 | AT4G18730 | AT5G45775 |
|---|---|---|---|---|
| AT2G42740 | 100 | 99.45 | 99.45 | 99.45 |
| AT3G58700 | 99.45 | 100 | 100 | 100 |
| AT4G18730 | 99.45 | 100 | 100 | 100 |
| AT5G45775 | 99.45 | 100 | 100 | 100 |

| RPL12 | AT2G37190 | AT3G53430 | AT5G60670 |
|---|---|---|---|
| AT2G37190 | 100 | 98.19 | 92.77 |
| AT3G53430 | 98.19 | 100 | 93.98 |
| AT5G60670 | 92.77 | 93.98 | 100 |

| RPS18 | AT4G09800 | AT1G34030 | AT1G22780 |
|---|---|---|---|
| AT4G09800 | 100 | 100 | 100 |
| AT1G34030 | 100 | 100 | 100 |
| AT1G22780 | 100 | 100 | 100 |

| RPL24 | AT2G44860 | AT2G36620 | AT3G53020 |
|---|---|---|---|
| AT2G44860 | 100 | 32.68 | 33.55 |
| AT2G36620 | 32.68 | 100 | 91.98 |
| AT3G53020 | 33.55 | 91.98 | 100 |

| RPS19 | At5g61170 | At3g02080 | At5g15520 |
|---|---|---|---|
| At5g61170 | 100 | 93.01 | 92.31 |
| At3g02080 | 93.01 | 100 | 95.8 |
| At5g15520 | 92.31 | 95.8 | 100 |

| RP used in Al-Jubran et al., 2013 | HMMR basic search (Arabidopsis thaliana) | Genes; Browning et al., 2015 |
|---|---|---|
| S18 | S19 | **At3g02080**, At5g15520, At5g61170 |
| S13 | S18 | At1g22780, At1g34030, **At4g09800** |
| L5 | L11 | At2g42740, At3g58700, **At4g18730**, At5g45775 |
| L11 | L12 | **At2g37190**, At3g53430, At5g60670 |
| S6 | S6 | **At4g31700**, At5g10360 |
| L24 | L24 | **At2g36620**, At3g53020, At2g44860 |
